# Longitudinal metabolomics data analysis informed by mechanistic models

**DOI:** 10.1101/2024.08.13.607724

**Authors:** Lu Li, Huub Hoefsloot, Barbara M. Bakker, David Horner, Morten A. Rasmussen, Age K. Smilde, Evrim Acar

## Abstract

**Motivation:** Metabolomics measurements are noisy, often characterized by a small sample size and missing entries. While data-driven methods have shown promise in terms of analyzing metabolomics data, e.g., revealing biomarkers of various phenotypes, metabolomics data analysis can significantly benefit from incorporating prior information about metabolic mechanisms. In this paper, we introduce a novel data analysis approach where data-driven methods are guided by prior information through joint analysis of simulated data generated using a human metabolic model and real metabolomics measurements.

**Results:** We arrange time-resolved metabolomics measurements of plasma samples collected during a meal challenge test from the COPSAC_2000_ cohort as a third-order tensor: *subjects* by *metabolites* by *time samples*. Simulated challenge test data generated using a human whole-body metabolic model is also arranged as a third-order tensor: *virtual subjects* by *metabolites* by *time samples*. Real and simulated data sets are coupled in the *metabolites* mode and jointly analyzed using coupled tensor factorizations to reveal the underlying patterns. Our experiments demonstrate that joint analysis of simulated and real data has a better performance in terms of pattern discovery achieving higher correlations with a BMI (body mass index)-related phenotype compared to the analysis of only real data in males while in females, the performance is comparable. We also demonstrate the advantages of such a joint analysis approach in the presence of incomplete measurements and its limitations in the presence of wrong prior information.

**Availability:** The code for joint analysis of real and simulated metabolomics data sets is released as a GitHub repository. Simulated data can also be accessed using the GitHub repo. Real measurements of plasma samples are not publicly available. Data may be shared by COPSAC through a collaboration agreement. Data access requests should be directed to Morten A. Rasmussen.

## Introduction

Human metabolism is a complex system, and deciphering this complex system is crucial in terms of understanding human health, diseases, and various phenotypes [36]. Metabolomics measurements of biological samples such as blood are rich sources of information, providing means to discover markers of various phenotypes, life style differences (e.g., diets, exercise), diseases, and reveal insights about the underlying metabolic mechanisms [33, 36]. Extensive biochemical knowledge including metabolic reactions is already available and has been compiled to construct computational models of human metabolism, e.g., Recon [47, 45]. These models have paved the way to whole-body models (WBM) constructed based on reactions driving the underlying molecular processes, the human anatomy and the physiology [46, 26]. However, there is still much to be unravelled to improve our understanding of the metabolism, and to achieve precision health [10].

A step towards deciphering this complex system has been to record the functioning of the metabolism using longitudinal measurements collected over time. For instance, at short time scales, time-resolved (dynamic) metabolomics data sets collected during meal challenge tests have been used to study the human metabolic response, linking observed differences to cardiometabolic diseases [27], and various phenotypes [53]. At longer time scales, metabolomics data, e.g., collected every few months, has shown promise in terms of revealing early signs of diseases, and the transition from healthy to diseased states [36, 37].

Analysis of such longitudinal metabolomics data is a challenging task due to high-dimensional, noisy and scarce/few measurements. Traditional methods rely on data summaries (e.g., data averaged across subjects [52], features summarizing time profiles using the area under the curve [13, 35]) or analysis of one feature at a time [12]. The workhorse data analysis methods in metabolomics remain univariate or multivariate methods [15] with analysis of variance [51, 13] and linear mixed models [32] commonly used in longitudinal metabolomics data analysis. Rather than relying on limited views of the data such as summary statistics or one feature at a time, recent studies have arranged time-resolved metabolomics measurements as a third-order tensor (also referred to as a three-way array) with modes: *subjects, metabolites, time*, and used tensor factorizations to reveal the underlying patterns, i.e., subject groups, metabolite clusters and temporal profiles [28, 53, 19, 41]. However, so far longitudinal metabolomics data analysis approaches have been limited to data-driven methods.

In this paper, we introduce a novel approach, where a data-driven method is informed by a computational model of metabolism. Recently, there have been significant efforts under various names (e.g., informed machine learning, physics-guided neural networks) to incorporate prior information in machine learning. These efforts mainly focus on supervised machine learning, and prior information in the form of simulations and knowledge graphs is usually integrated at the training data stage through data augmentation or used to constrain deep neural networks by simplifying architectures or penalizing the loss functions [48]. Here, instead we incorporate mechanistic models in unsupervised learning through a novel data fusion approach that jointly analyzes simulated data generated using a mechanistic model and real data. In particular, we focus on the analysis of time-resolved metabolomics measurements of plasma samples collected during a meal challenge test. We arrange the real data as a third-order tensor: *subjects, metabolites, time samples*. Simulated challenge test data generated using a human whole-body metabolic model is also arranged as a third-order tensor: *virtual subjects, metabolites, time samples*. Real and simulated data sets, which are coupled in the *metabolites* mode, are then jointly analyzed using coupled tensor factorizations (See Figure 1). We demonstrate that by guiding the analysis of noisy real data using clean simulated data, the proposed joint analysis approach achieves improved performance in terms of pattern discovery revealing patterns with higher correlations with a BMI (body mass index)-related phenotype compared to the analysis of only real data. We also demonstrate the advantages of incorporating prior information using such a data fusion approach in the presence of incomplete measurements, and discuss the limitations in the presence of wrong prior information.

**Fig. 1.**
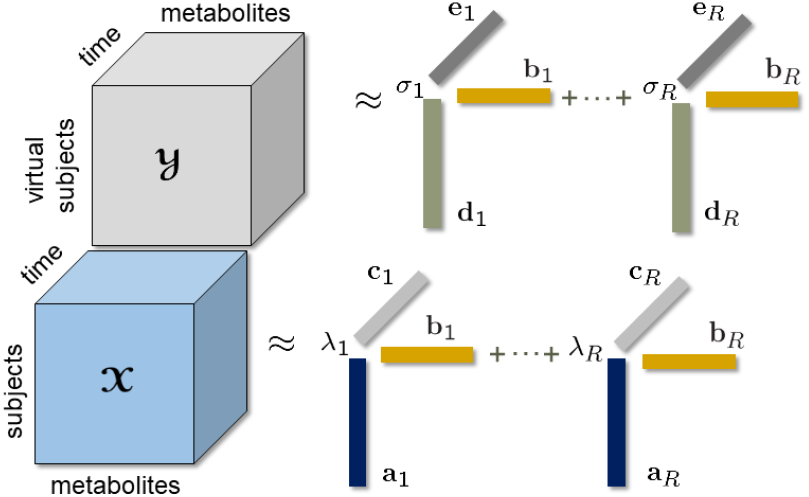
An *R*-component ACMTF model jointly analyzing third-order tensors 𝒳 (*subjects* by *metabolites* by *time*) and 𝒴 (*virtual subjects* by *metabolites* by *time*) coupled in the *metabolites* mode.

## Materials and Methods

### Real Meal Challenge Test Data

The real data corresponds to measurements of specific hormones and Nuclear Magnetic Resonance (NMR) spectroscopy measurements of blood samples collected during a meal challenge test from the COPSAC_2000_ cohort [14]. The cohort consists of 411 healthy subjects (with mothers with a history of asthma). The data in this paper comes from 299 of those generally healthy subjects who underwent a meal challenge test at the age of 18. Blood samples were collected from the participants after overnight fasting and also following a standardized mixed meal [44]. The meal was a hot beverage consisting of palm oil, glucose, and skimmed milk powder.

Blood samples were collected at 15, 30, 60, 90, 120, 150 and 240 minutes after the meal intake. The COPSAC_2000_ study was conducted in accordance with the Declaration of Helsinki and was approved by the Copenhagen Ethics Committee (KF 01-289/96 and H-16039498) and the Danish Data Protection Agency (2015-41-3696). The study participants gave written consent.

Plasma samples were then measured using NMR through the Nightingale Blood Biomarker Analysis, which provides 250 features for each sample. Features include lipoproteins, apolipoproteins, amino acids, fatty acids, glycolysis-related metabolites, ketone bodies and an inflammation marker. Details about the meal challenge test, sample preparation and the full list of features are given in [53]. For each participant, additional meta information including body composition measures and HOMA-IR (Homeostatic model assessment for Insulin Resistance, an insulin resistance measure) is also available. Descriptive statistics of these meta variables (i.e., HOMA-IR, weight, height, waist circumference, BMI, waist/height ratio, muscle mass, fat mass, body fat percentage, muscle to fat ratio, fat mass index, and fat free mass index) stratified by sex are given in [53].

To investigate metabolic differences among subjects in response to a meal challenge, we previously analyzed both fasting and T0-corrected data (where postprandial data is corrected by subtracting the fasting state measurements) for this cohort. Our analysis revealed static and dynamic biomarkers of a BMI-related phenotype as well as gender-related differences [53, 30]. In this study, we consider six features out of the complete set of measurements since these six features are the common blood metabolites in the WBM model, including insulin (Ins), glucose (Glc), pyruvate (Pyr), lactate (Lac), alanine (Ala) and *β*-hydroxybutyrate (Bhb). We focus on analyzing T0-corrected data, as previous studies have demonstrated its effectiveness in terms of capturing dynamic biomarkers [29, 53]. Six subjects (three males and three females) were removed before the analysis since two subjects had a large amount of missing data, and four subjects had extreme levels of acetate (greater than 0.4 mmol/l) probably due to a recent alcohol exposure. Measurements are arranged as a third-order tensor with modes: *subjects, metabolites, time samples*. The tensor is of size 141 subjects × 6 metabolites × 7 time samples for males, and 152 subjects × 6 features × 7 time samples for females.

### Simulated Meal Challenge Test Data

We generate simulated postprandial metabolomics data using a human whole-body metabolic model, which involves 202 metabolites, 217 reaction rates, and 1140 kinetic parameters [26]. The model is based on ordinary differential equations, capturing the complexity of multi-organ interactions and incorporating insulin and glucagon regulation after a meal intake. In this model, a meal containing 87 grams (g) carbohydrate and 33 g fat is considered after a 10-hour fasting. Note that the simulated meal composition differs from that of the real meal, but the responses of key glycolysis-related metabolites to the meal challenge are in the physiological range. A detailed comparison of the real and simulated meal as well as the time profiles following the meal challenge is available in [29]. For each subject, the data is generated as follows: First, 10-hour fasting concentrations are acquired for each individual by running the human WBM model with a unique set of randomly perturbed kinetic constants in the liver. We use the default initial values for model variables from [26], but specific adjustments are made for initial concentrations of certain blood metabolites, i.e., insulin, glucose, pyruvate, lactate, alanine, *β*-hydroxybutyrate, triglyceride and total cholesterol are set to the median of fasting concentrations in the real data. Subsequently, the simulation advances to the meal challenge phase, and runs the human WBM model using the 10-hour fasting state of each individual as initial values. Concentrations of metabolites are recorded at specific time points (aligned with the measurements in real data). The metabolic model provides simulated concentrations of 202 metabolites from blood and eight different organs. In this study, we use concentrations of six blood metabolites, i.e., Ins, Glc, Pyr, Lac, Ala, and Bhb, that are also measured in real data.

We generate 50 virtual control subjects without introducing any group differences, and with individual variations introduced through random perturbations of the kinetic parameters in the liver. For each kinetic parameter, a random perturbation of up to 20% of its default value is introduced^1^ (See [29] for more details on individual variations). We have observed that a sample size of around 50 or more subjects is needed in order to extract robust patterns. With less number of subjects, e.g., 10 subjects, only idiosyncratic behavior is captured. The simulated T0-corrected metabolomics data is arranged as a third-order tensor of size 50 subjects × 6 metabolites × 7 time samples. Note that since our goal is to guide the analysis of real data through clean patterns extracted from virtual subjects, and the real subjects in the cohort are healthy, when generating data for virtual subjects, we keep the individual variation low and do not introduce any patterns that would be expected in diseases.

### Tensor Factorizations

As an extension of matrix factorizations to higher-order data sets (also known as multiway arrays), tensor factorizations are used to extract the underlying patterns in multiway arrays [7, 25]. They have been successfully used in many domains including neuroscience [1, 50], chemometrics [42], and social network analysis [34]. Among different tensor factorization methods, we focus on the CANDECOMP/PARAFAC (CP) tensor model [17, 20], also known as the canonical polyadic decomposition [21] to analyze time-resolved metabolomics data sets. The CP model extracts the underlying patterns uniquely under mild conditions [25]. Uniqueness properties of the CP model facilitate interpretation, which is particularly important when the goal is to discover phenotypes and biomarkers in metabolomics data analysis.

Given a third-order tensor 𝒳 ∈ ℝ^*I*×*J*×*K*^, an *R*-component CP model represents the data as the sum of minimum number of rank-one tensors as follows:

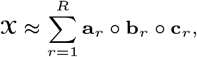

where ∘ denotes the vector outer product, and **a**_*r*_, **b**_*r*_ and **c**_*r*_ correspond to the *r*th column of factor matrices **A** ∈ ℝ^*I*×*R*^, **B** ∈ ℝ^*J*×*R*^ and **C** ∈ ℝ^*K*×*R*^, respectively. The CP model can also be denoted as 𝒳 ≈ ⟦***λ***; **A, B, C**⟧. ***λ*** ∈ ℝ^*R*^ can absorb the weights of the rank-one tensors, i.e., **a**_*r*_ ∘ **b**_*r*_ ∘ **c**_*r*_ for *r* = 1, …, *R*, by normalizing columns of the factor matrices to unit norm. The rank-one components reveal the underlying patterns in the data, e.g., if 𝒳 is a *subjects* by *metabolites* by *time samples* tensor, the *r*th CP component may reveal subject stratifications in **a**_*r*_, groups of metabolites responsible for the stratification in **b**_*r*_, and their temporal pattern in **c**_*r*_. The promise of the CP model in terms of revealing subject stratifications/phenotypes and underlying metabolic mechanisms from time-resolved metabolomics data has been demonstrated in recent studies [29, 53]. The CP model is unique up to permutation and scaling ambiguities under mild conditions, where the permutation ambiguity indicates that the order of rank-one tensors is arbitrary, and the scaling ambiguity corresponds to arbitrarily scaling the vectors in each rank-one tensor as long as the product of the norms stays the same. Since these ambiguities do not interfere with the interpretation of the extracted patterns, the CP model has been widely used in many applications, e.g., longitudinal microbiome data analysis [31], analysis of neuroimaging signals [1, 22], when the goal is to interpret the underlying patterns to extract insights from complex data. The model is often fit to the data by solving the following optimization problem:

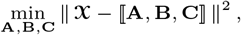

where ∥ ∥ denotes the Frobenius norm for matrices and higher-order tensors, and 2-norm for vectors. The Tikhonov regularization can be included by adding *γ*(∥ **A** ∥^2^ + ∥ **B** ∥^2^ + ∥ **C** ∥^2^) to the objective function.

### Coupled Tensor Factorizations

In this paper, we consider incorporating the prior knowledge encapsulated in computational models of human metabolism by jointly analyzing simulated data (in the form of tensor 𝒴 of size *I*_virtual_ by *J*_virtual_ by *K*_virtual_ with modes: *virtual subjects, metabolites, time samples*) generated using a human WBM model, and the real data (in the form of tensor 𝒳 of size *I*_real_ by *J*_real_ by *K*_real_ with modes: *subjects, metabolites, time samples*). We focus on the case where 𝒳 and 𝒴 are coupled in the *metabolites* mode as in Figure 1 with *J*_virtual_ = *J*_real_ = *J*, i.e., only common metabolites in simulated and real data are taken into account in the analysis.

An effective approach to jointly analyze such higher-order data sets is to use coupled tensor factorizations [2, 4, 34, 43]. Given tensors 𝒳 and 𝒴 coupled in the *metabolites* mode, a coupled CP model jointly analyzes them by modeling each tensor using a CP model and extracting the same factor matrix from the coupled mode, e.g., the same **B** factor matrix in both CP models as in Figure 1. In particular, we use the structure-revealing CMTF model (also known as the advanced Coupled Matrix and Tensor Factorizations (ACMTF)) [5], which jointly analyzes each data set while trying to learn shared and unshared factors.

Given 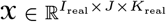 and 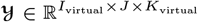 coupled in the second mode, e.g., *metabolites* mode, an *R*-component ACMTF model jointly analyzes these data sets by solving the following optimization problem:

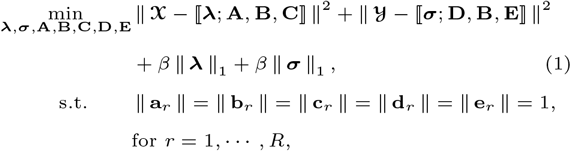

where columns of factor matrices 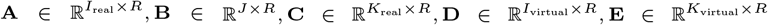 are constrained to be unit norm, i.e., ∥ **a**_*r*_ ∥ = 1, ∥ **b**_*r*_ ∥ = 1, ∥ **c**_*r*_ ∥ = 1, ∥ **d**_*r*_ ∥ = 1, ∥ **e**_*r*_ ∥ = 1 for *r* = 1, …, *R*. With normalized factor vectors in each mode, ***λ*** ∈ ℝ^*R*^ and ***σ*** ∈ ℝ^*R*^ contain weights of the rank-one terms in each data set. ∥ · ∥_1_ denotes 1-norm of a vector. By enforcing sparsity on the weights through the 1-norm penalty, when *β >* 0, the ACMTF model tries to reveal unshared factors with zero or close to zero weights. Coupled CP models inherit uniqueness properties from the CP model [43], and sparsity penalties on the weights have been effective to reveal shared and unshared factors [5]. Note that unit norm constraints already act as regularization; therefore, we do not consider further regularization of the model.

Other types of coupling between real and simulated data sets are also possible. For instance, data sets can be coupling in the *time* mode. However, uncoupled time patterns may facilitate the discovery of discrepancies between simulations and real data as we observe in the experiments. Data sets can also be coupled in both *metabolites* and *time* modes through concatenation in *subjects* mode and analyzed as a single tensor, for instance using a CP model. In such a setting, the model would focus on modelling the difference between virtual and real subjects rather than focusing on stratifications among real subjects. Therefore, in this paper, we consider coupling the data sets only in *metabolites* mode.

CMTF models have been used in many domains, e.g., social network analysis [34], remote sensing [23], neuroscience [6, 22], chemometrics [2] and metabolomics [30]. However, to the best of our knowledge, this is the first application of such a model to incorporate mechanistic models in unsupervised data mining.

### Experimental set-up

#### Data Preprocessing

Before the analysis, third-order tensors (𝒳 and 𝒴) representing T0-corrected metabolomics data with modes *subjects* by *metabolites* by *time samples* are preprocessed by centering across the *subjects* mode to remove mean-based offsets and then scaling within the *metabolites* mode (i.e., each metabolite slice is divided by the root mean squared value of that slice) to ensure similar scales across all metabolites. See [16] for centering and scaling of higher-order tensors. After preprocessing, when fitting the ACMTF model, each tensor is divided by its Frobenius norm to give equal importance to each data set in (1).

#### Implementation Details

We use `acmtf_opt` from the CMTF toolbox^2^ to fit the ACMTF model [5], and `cp_wopt` [3] from the Tensor Toolbox (version 3.1) [11] to fit the CP model. The nonlinear conjugate gradient algorithm from the Poblano Toolbox [18] is used to solve the optimization problems for CP and ACMTF. In the case of missing entries, functions `acmtf_opt` and `cp_wopt` use weighted optimization fitting the model only to the known entries. The sparsity penalty parameter *β* in (1) is set to 10^*−*3^ [5]. Multiple random initializations are used to avoid local minima when fitting ACMTF and CP models, and the initialization that yields the lowest function value is chosen as the best run for further analysis. All experiments are conducted using MATLAB 2020a. For more details, see the GitHub repository ^3^.

#### Model Selection

Selecting the right number of components (*R*) is crucial for extracting the underlying patterns accurately for CP and ACMTF models. Here, we rely on the replicability of the extracted patterns across random subsets of the real subjects [8, 53]. The replicability is assessed by fitting the model (CP or ACMTF) to subsets of subjects, where 10% of subjects from the real data 𝒳 in Figure 1 is randomly removed. The similarity of the patterns extracted from different subsets of subjects (after finding the best matching permutation) is computed using the Factor Match Score (FMS). FMS_𝒳_ and FMS_𝒴_ are defined as:

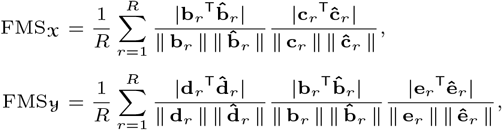

where ⟨**b**_*r*_, **c**_*r*_, **d**_*r*_, **e**_*r*_⟩ and 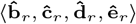 are the *r*th column of factor matrices from *R*-component ACMTF models in the *metabolites* (coupled mode), *time* (real), *subjects* (virtual) and *time* (virtual) modes. When assessing the replicability of CP models, only FMS_𝒳_ is considered. FMS values range between 0 and 1, where 1 indicates an exact match between the components of models fitted to different subsets of subjects.

The *model fit* is also computed to determine how well the models explains the data sets. The fit is defined as:

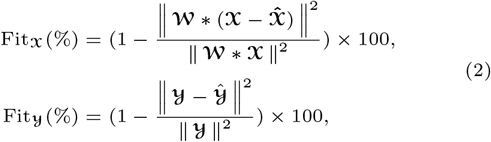

where 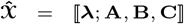 and 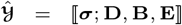 are the approximations of real and simulated data, respectively. The binary tensor 𝒲 indicates observed (*w*_*ijk*_ = 1) or missing (*w*_*ijk*_ = 0) entries in tensor 𝒳. The symbol * denotes the (element-wise) Hadamard product. A fit value close to 100% implies that the model explains the data well.

## Results

Previously, gender differences were observed in the COPSAC_2000_ cohort in terms of how the dynamic metabolic response was related to a BMI-related group difference [53]. Therefore, in this study, we analyze males and females separately. In males, we demonstrate that joint analysis of simulated and real metabolomics data has a better performance in terms of pattern discovery achieving higher correlations with a BMI-related phenotype compared to the analysis of only real data. In females, joint analysis of simulated and real data sets achieves similar performance compared to the analysis of only real data. Using a larger set of metabolites from the same meal challenge study, we previously observed that correlations were much lower for females possibly due to anthropometric differences (e.g., where fat is deposited in males vs. females) [53]. Here, considering a specific set of metabolites, higher correlations are achieved in females -however, they are still much lower than males. As the improvement may be limited due to anthropometric differences and the WBM model validation mainly relies on data from males [26], in the rest of the paper, we focus on the analysis of measurements from males, and discuss the results for females (given in Supplemental File (Section A)) in the Discussion section.

We evaluate the performance of the methods in terms of how well they reveal a BMI-related phenotype characterized by the meta variables. More specifically, the performance is assessed in terms of the correlations between subject scores (captured by the methods) and BMI-related meta variables. Note that as the subjects in the cohort are generally healthy, there is no variation due to any specific disease that is expected to be picked up by the methods.

### Analysis of real metabolomics data

We analyze the real T0-corrected metabolomics measurements from males 𝒳 (141 males × 6 metabolites × 7 time points) using a 3-component CP model with Tikhonov regularization (with regularization parameter *γ* = 0.01). See Supplemental File (Section B.1) for the selection of number of components and the regularization parameter. The model explains 52.4% of the data. Figure 2 shows the factors extracted by the 3-component CP model.

**Fig. 2.**
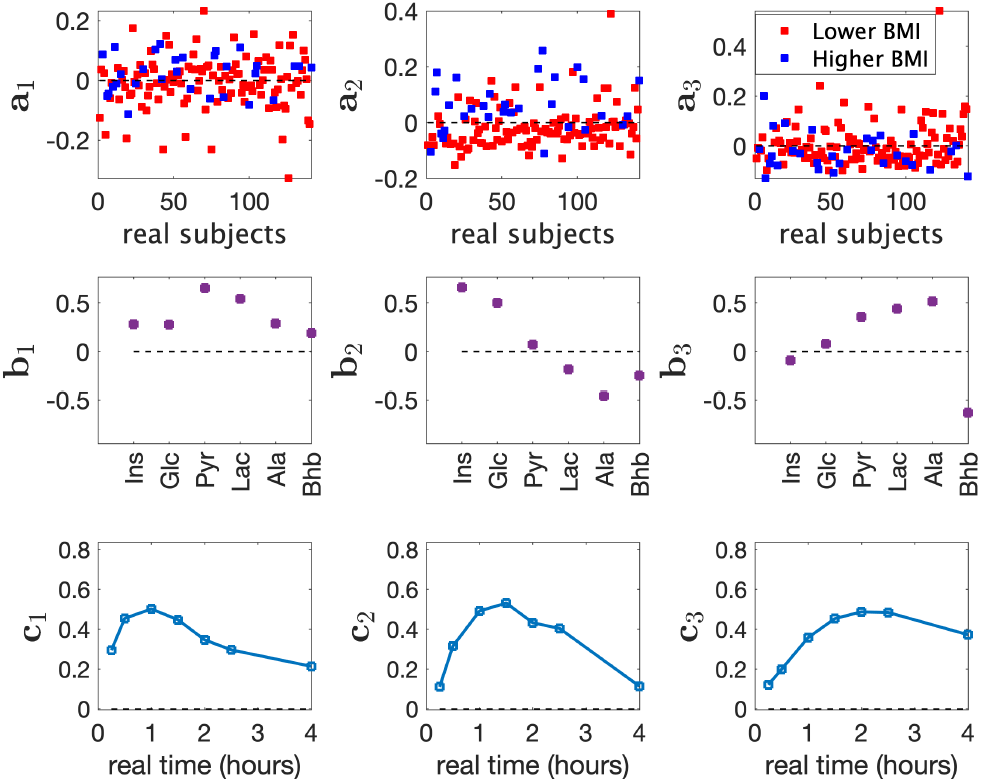
Factors of the 3-component CP model of T0-corrected data from males. ⟨**a**_*r*_, **b**_*r*_, **c**_*r*_ ⟩, *r* = 1, 2, 3, are the components in the *subjects, metabolites* and *time* modes.

We observe that the model reveals a weak BMI-related group difference in the second component (*p*-value = 1 × 10^*−*4^ using a two-sample *t*-test on **a**_2_ in Figure 2), where *Lower BMI* and *Higher BMI* corresponds to BMI *<* 25 and BMI ≥ 25, respectively. In the metabolites mode (**b**_2_), Ins, Glc and Ala have the largest score values (in terms of absolute value) indicating that they are the most related metabolites to BMI group difference. In particular, Ins and Glc have positive values indicating that changes in these metabolites are positively related to *Higher BMI* while the change in Ala is negatively related to *Higher BMI*. In the time mode, **c**_2_ increases until around 1.5 hours and decreases afterwards showing the temporal profile of the metabolic response modelled by this component. Although we discuss subject group differences based on BMI groups, **a**_2_ is also correlated with other meta variables as shown in Figure 4a.

The first component ⟨**a**_1_, **b**_1_, **c**_1_⟩ and the third component ⟨**a**_3_, **b**_3_, **c**_3_⟩ in Figure 2 potentially model non-BMI related individual differences in the data. The first component mainly models an early response captured by **c**_1_, and the third component models a late response captured by **c**_3_. The third metabolite factor, i.e., **b**_3_, reveals that changes in Pyr, Lac and Ala behave opposite to the change in Bhb, which aligns with the observation that the concentrations of Pyr, Lac and Ala increase while Bhb decreases after the meal intake as shown in the temporal profiles of these metabolites in Figure S.5 in Supplemental File. No statistically significant BMI-related group difference is observed in these components, and correlations between subject scores and meta variables are less than or around 0.2 (except for HOMA-IR -for which the third component reveals a correlation of 0.32).

### Joint analysis of real and simulated metabolomics data

We jointly analyze the real T0-corrected metabolomics data 𝒳 (141 males × 6 metabolites × 7 time points) and simulated metabolomics data 𝒴 (50 subjects × 6 metabolites × 7 time points) using a 3-component ACMTF model by coupling the data sets in the *metabolites* mode. The model explains 50.0% of the real data, and 71.8% of the simulated data. See Supplemental File (Section B.2) for the selection of the number of components.

Figure 3 shows the factors extracted using a 3-component ACMTF model. The second component (**a**_2_) reveals a BMI-related group difference (*p*-value = 3×10^*−*6^). In the metabolites mode **b**_2_, Ins and Glc have the largest absolute score values showing positive association with *Higher BMI*. The first and third components may be modelling non-BMI related individual variations in the data. No statistically significant BMI-related group difference is observed in these components and correlations with meta variables are all much smaller than 0.2, with largest ones around 0.2. These components have similar dynamic patterns in the real data part (i.e., **c**_1_ and **c**_3_); however, corresponding metabolite factors, **b**_1_ and **b**_3_, model the behavior of different metabolites, i.e., Pyr, Lac and Ala have large score values on **b**_1_ while Bhb is mainly modelled by **b**_3_.

**Fig. 3.**
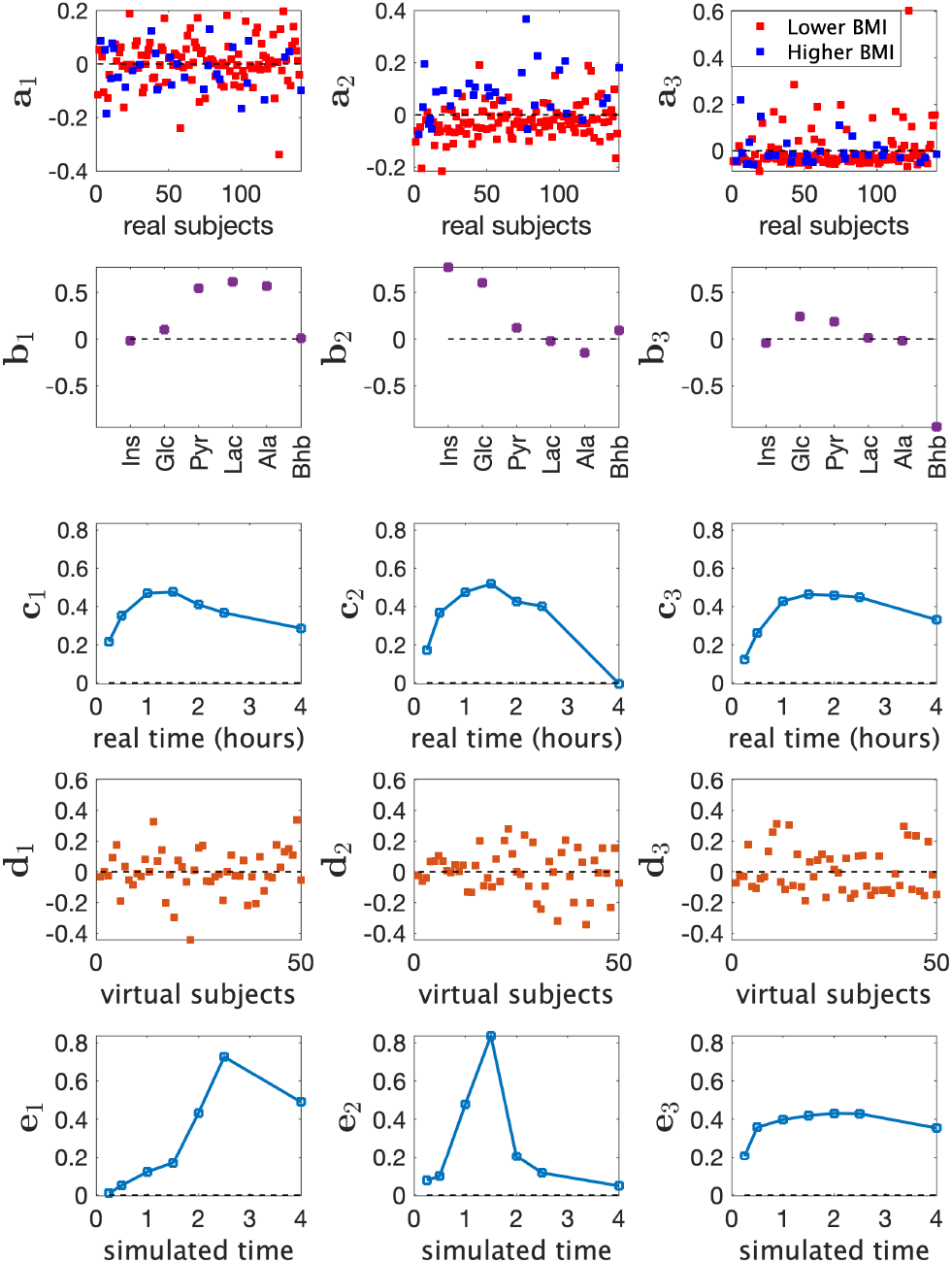
Factors of the 3-component ACMTF model of T0-corrected real data (from males) and simulated data. ⟨**a**_*r*_, **b**_*r*_, **c**_*r*_, **d**_*r*_, **e**_*r*_ ⟩, *r* = 1, 2, 3, are the components in the *subjects* (real), *metabolites* (coupled mode), *time* (real), *subjects* (virtual) and *time* (virtual) modes.

We do not observe any apparent clustering among the virtual subjects (**d**_1_, **d**_2_ and **d**_3_) since no group-specific information was incorporated during the generation of the simulated data. Consequently, virtual subject patterns mainly reflect individual variations in the simulated data.

As the data sets are coupled only in the metabolites mode through an ACMTF model, the model reveals time profiles specific to each data set. When time profiles **c**_1_, **c**_2_, **c**_3_ extracted from the real data are compared with the ones from the simulated data, i.e., **e**_1_, **e**_2_, **e**_3_, we observe that the model captures different temporal profiles from real and simulated data potentially revealing the discrepancies between simulated and real data. This discrepancy may be due to the fact that virtual and real subjects undergo meal challenges with different contents, and this may result in the observed differences in temporal profiles as discussed in [29].

### Analysis of real data vs. joint analysis of simulated and real data

Figure 2 vs. Figure 3 show that joint analysis of simulated and real data reveals cleaner patterns in the metabolites mode affected less by the noise in real data: (i) Pyr, Lac and Ala are close to each other with large score values in both **b**_1_ and **b**_3_ in Figure 2. On the other hand, they are mainly modelled by **b**_1_ using the joint analysis as shown in Figure 3; (ii) Ins and Glc cluster closely with large score values in both **b**_1_ and **b**_2_ in Figure 2 while they are modelled mainly by the second factor (**b**_2_ in Figure 3) in the joint analysis; (iii) Bhb has contributions in all components in Figure 2 whereas in Figure 3 Bhb only contributes to **b**_3_. As a result of these differences, we observe that the BMI-related component captured through the joint analysis of simulated and real data, i.e., **b**_2_ in Figure 3, shows higher correlations with all meta variables as shown in Figure 4a.

Another observation is that the CP analysis of real data reveals a potential negative association between Ala and *Higher BMI* group (as shown in ⟨**a**_2_, **b**_2_, **c**_2_⟩ in Figure 2). This is a pattern not evident in the joint analysis. Time profiles of raw data including Ala are given in Figure S.5 of Supplemental File. These plots show that there are differences in Ala concentrations between the *Higher* and *Lower BMI* groups (which are statistically significant at some time points). The joint analysis does not identify Ala as an important metabolite related to BMI group difference since the simulated data does not support such a pattern. For virtual subjects, we observe different temporal profiles for Ala compared to Ins, Glc and Bhb, and similar temporal profiles compared to Pyr and Lac. This prevents the joint analysis to extract a metabolite pattern similar to **b**_2_ in Figure 2, and facilitates the extraction of **b**_1_ and **b**_2_ in Figure 3. For the CP analysis of only simulated data, see Figure S.6a in Supplemental File.

### Joint analysis of real and simulated metabolomics data in the presence of missing data

Missing data may be observed in metabolomics data analysis due to various reasons such preprocessing issues of the raw metabolomics measurements or sample handling problems. These errors may cause random missing measurements or missing measurements for a whole sample. We demonstrate that joint analysis of incomplete real data with simulated data can improve the pattern discovery performance. Here, we randomly set 10% of the real data to be missing, including 5% of the data corresponding to missing fibers. Here, a fiber corresponds to measurements of all metabolites from a sample, i.e., from a specific subject at a certain time point. The remaining 5% corresponds to randomly missing entries (i.e., a single measurement). The incomplete real measurements are then analyzed using a CP model and also jointly analyzed with simulated data using an ACMTF model. We generate and analyze 32 such randomly incomplete data sets. Figure 4b reports the correlations (of the subject scores from the BMI-related component) with meta variables when using an ACMTF model vs. a CP model. Boxplots contain the correlations from the analysis of 32 data sets. Figure 4b shows that joint analysis demonstrates more consistent and higher correlations than the CP model.

**Fig. 4.**
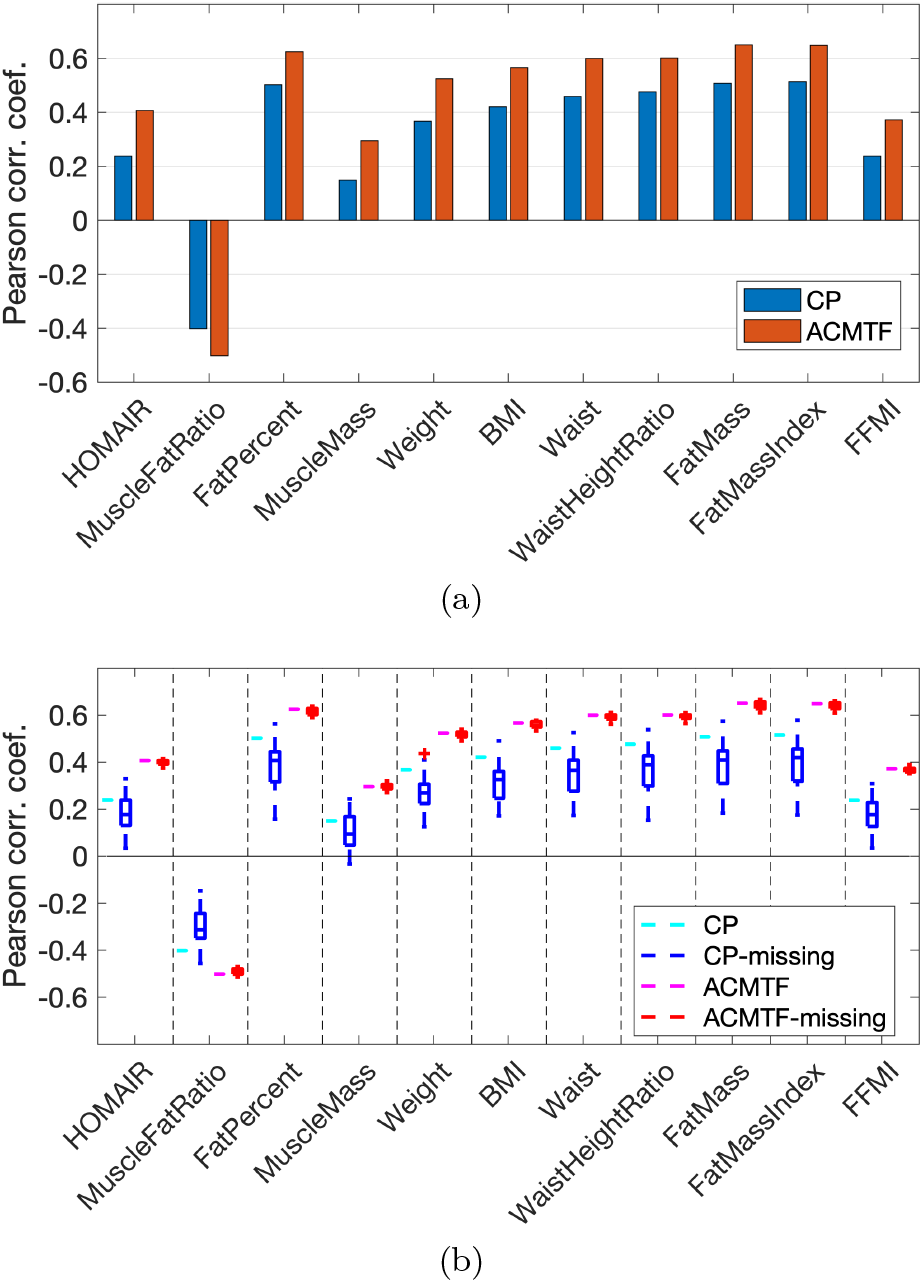
Correlations between subject scores and meta variables for the factor showing BMI-related group difference in CP and ACMTF models using (a) the real T0-corrected data 𝒳 from males. Here, values correspond to the correlations between **a**_2_ in Figure 2 and meta variables for CP, and between **a**_2_ in Figure 3 and meta variables for ACMTF, (b) incomplete real T0-corrected data from males, where 10% of the entries in 𝒳 is set to missing. 32 randomly incomplete data sets are considered. Correlations achieved using CP and ACMTF models of the original real data are included again in (b) for easier comparison. Meta variables correspond to HOMAIR: Homeostatic model assessment for Insulin Resistance; MuscleFatRatio: Muscle to fat ratio; FatPercent: Body fat percentage; MuscleMass: Amount of muscle in the body (kg); Weight: Weight (kg); BMI: Body Mass Index; Waist: Waist circumferance (cm); WaistHeightRatio: Waist measurement divided by height (cm); FatMass: Amount of body fat (kg); FatMassIndex: FatMass divided by height^2^; FFMI: Fat Free Mass Index.

### Joint analysis of real and simulated metabolomics data in the presence of conflicting information

While we have demonstrated the effectiveness of joint analysis of real and simulated data, it is important to note that it is possible to have conflicting information between the prior information (e.g., simulated data) and real data. Such conflicting information may prevent revealing the underlying patterns accurately. To demonstrate the performance of joint analysis in the presence of conflicting information, we create simulated data with wrong prior information as follows:

Step 1. *Default pattern extraction and residual calculation*. We use a 3-component CP model to extract the underlying patterns from the simulated T0-corrected data (see Figure S.6a in Supplemental File). The data approximated by the model is denoted by 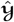, and residuals by ε,

Step 2. *Conflicting pattern construction*. The first and third components (from Step 1) are retained while the second component is modified by introducing wrong prior information. In the default (correct) pattern, Ins and Glc are close to each other having large positive values in the second component while values of the remaining metabolites are close to zero. We break down the positive association between Ins and Glc to introduce conflicting information. This is wrong prior information for the real data since the cohort consists of healthy subjects where such relation is not expected to be observed. Loading values of Ins, Glc, Pyr, Lac, Ala, Bhb are set to 1, −1, 0, 0, 0, 0, respectively, and the factor vector is normalized (i.e., divided by its 2-norm). See Figure S.6b in Supplemental File for the modified pattern.

Step 3. *Construction of simulated data with conflicting information*. Tensor 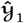 is then constructed using the modified CP patterns. The simulated data with conflicting information, denoted by 𝒴_1_, is obtained by adding the residual term (obtained in Step 1) to 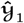, i.e., 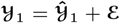.

Figure 5 shows that coupling with conflicting prior information leads to worse correlations between subject scores and meta variables compared to the analysis of real data using a CP model. Such poor correlations stem from the fact that the joint analysis obstructs the extraction of correct patterns from the real data. This issue occurs because these correct patterns, in particular the second component in Figure 3 mainly modelling Ins and Glc, are substantially different from the ‘broken-down’ pattern, i.e., the second component in Figure S.6b in Supplemental File, in the simulated data. We observe that the ACMTF model extracts the ‘broken-down’ pattern (see Figure S.7 in Supplemental File). Here, we also observe that the ACMTF model explains less of the real data, and more of the simulated data compared to the case when the real data is jointly modelled with the default simulated data. In other words, the model fit of the real data part drops from 50.0% to 44.0% while the model fit of the simulated data increases from 71.8% to 74.3% when the default simulated data is replaced with the simulated data containing conflicting information in the joint analysis. When we look at weights of the components (i.e., *λ, σ* in Figure 1) learned by ACMTF models given in Figure 6a and 6b, we observe a decrease in the weight of the second component (Ins-Glc related pattern) in the real data part (i.e., *λ*_2_) while *λ*_1_ and *λ*_3_ remain relatively unchanged - which is consistent with the observed decrease in the model fit. Although we observe a decrease in *λ*_2_, the second component still looks like a shared pattern. *λ*_2_ close to 0 in this case would indicate an unshared factor and would make the identification of conflicting information possible. This shows the limitation of the ACMTF model in terms of detecting conflicting information in the case of noisy data sets.

**Fig. 5.**
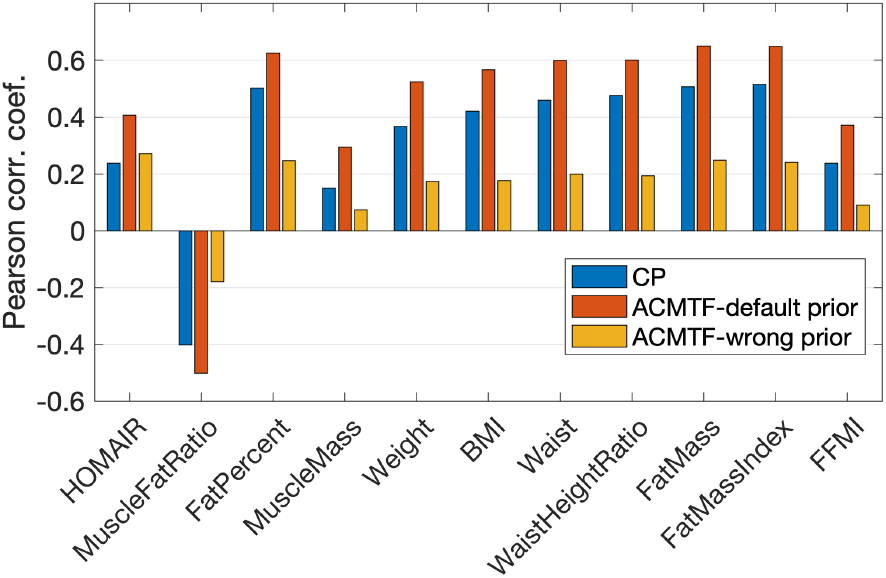
Correlations between the subject scores and meta variables for the factor showing BMI-related group difference - captured using the CP model of T0-corrected real data, ACMTF model of T0-corrected real data and the default simulated data, and ACMTF model of T0-corrected real data and simulated data with conflicting information.

**Fig. 6.**
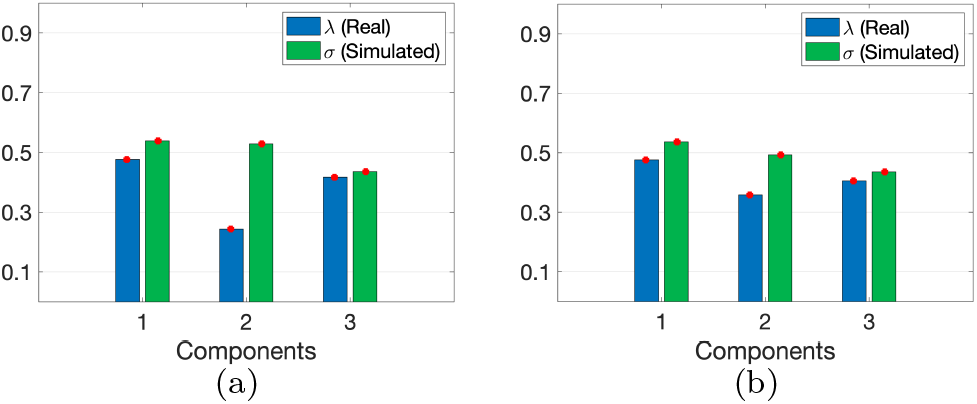
Weights of the components in ACMTF models of (a) T0-corrected real data and wrong simulated data, (b) T0-corrected real data with default simulated data.

## Discussion

Longitudinal metabolomics measurements collected over time hold the promise to improve our understanding of the metabolism, reveal early signs of diseases and facilitate precision health. Recent technological advancements have facilitated the collection of such time-resolved metabolomics measurements [39]. However, analysis of such data sets has many challenges including noisy, missing measurements, and small sample size. In this paper, we have introduced a novel data analysis approach for longitudinal metabolomics data by incorporating mechanistic models based on prior biological knowledge in order to guide the analysis of noisy real data with clean prior information. We jointly analyze time-resolved metabolomics measurements and simulated data generated using a human WBM model. Our experiments demonstrate that the proposed approach achieves better pattern discovery performance compared to the analysis of only real data. A similar performance improvement has also been demonstrated in the presence of real data with missing entries. This enhanced performance is attributed to the extraction of cleaner patterns facilitated by joint analysis of real data with clean simulated data.

Compared to the improved performance in males (in terms of correlations between subject scores extracted by the models and various BMI associated variables), in females informing the real data analysis with simulated data through joint analysis of real and simulated metabolomics measurements does not change the performance much as shown in Figure S.1 in Supplemental File. In both males and females, we observe that joint analysis of real and simulated data reveals a component in the metabolite mode mainly modelling Ins and Glc (see **b**_2_ in Figure S.2 in Supplemental File) due to the existence of such a pattern in simulated data (see **b**_2_ in Figure S.6a) and that is the component revealing BMI-related group difference. As the Ins/Glc-centric component is more tightly associated with the BMI-associated variables compared to Lac, Ala, BhB modelled in that component using the CP model of the data from males, having a cleaner component focusing on Ins/Glc improves the correlations with meta variables for males. However, in females, that component is already dominated by Ins/Glc in the CP model. Therefore, joint analysis does not change the subject scores for females much and the correlations stay almost the same.

When jointly analyzing real and simulated data sets, we give equal importance to each data set. Determining the optimal weights in coupled factorizations remains an open research question [49, 40, 38]. We assess the sensitivity of joint analysis to different weighting schemes and observe that unless we are close to the extremes (i.e., only modelling the real data or only modelling the simulated data), the ACMTF model is not very sensitive to the weight selection (see Section E of Supplemental File). While this finding is promising, it is still not conclusive in terms of determining the optimal weights. Nonetheless, it supports assigning equal weights to real and simulated data in the absence of prior information.

Potential discrepancies between real and simulated data sets may arise due to, for instance, incomplete knowledge about the underlying system. In our experiments, we have demonstrated that while clean prior information improves pattern discovery performance, the performance of the proposed data fusion approach degrades in the presence of wrong prior information. Therefore, it is crucial to investigate methods for detecting such conflicting information, necessitating robust diagnostic tools for detecting shared and unshared factors across data sets [5, 24]. Revealing shared and unshared factors could potentially uncover new mechanisms or identify erroneous information in computational models. Consequently, such advancements would not only enhance the analysis of real data but also facilitate the development of alternative methods for validating computational models, and improved understanding of deviations in computational models from reality [9].

As future work, we also plan to focus on different settings in simulations, and study how they affect the performance of real data analysis. In particular, we will consider different numbers of virtual subjects to account for stronger/weaker prior information, and different levels of individual variation in the simulations. We expect that these settings will play a role in both pattern discovery and model selection using the replicability test. Furthermore, as seen in our experiments, simulated and real metabolomics often have partially overlapping sets of metabolites. In the experiments we have focused on only the matching metabolites. We plan to consider different types of coupling between real and simulated data sets in order to incorporate all metabolite measurements. Recent modelling and algorithmic advances in coupled tensor factorizations enable joint analysis of data sets with such coupling relations [38].

## Author contributions statement

E.A. and L.L. conceived the experiments; L.L. conducted the experiments; L.L., H. H., A.K.S., E. A., D.H., B.M.B., M.A.R. analyzed the results; L.L. and E.A. wrote the manuscript. All authors reviewed the manuscript.

## Acknowledgments

We thank the children and families of the COPSAC_2000_ cohort for their contribution, and the clinical team at COPSAC for conducting the clinical study. This work was supported in part by the Research Council of Norway through project 300489 and in part by the Novo Nordisk Foundation grant NNF19OC0057934.

## Footnotes

1 The simulated data has been released in a GitHub repository together with [29]: https://github.com/Lu-source/project-of-challenge-test-data/

2 https://github.com/eacarat/CMTF_Toolbox

3 https://github.com/Lu-source/ACMTF_Real_Simulated

## References

1. E. Acar, C. A. Bingol, H. Bingol, R. Bro, and B. Yener. Multiway analysis of epilepsy tensors. Bioinformatics, 23(13):i10–i18, 2007.

2. E. Acar, R. Bro, and A. K. Smilde. Data fusion in metabolomics using coupled matrix and tensor factorizations. Proceedings of the IEEE, 103:1602–1620, 2015.

3. E. Acar, D. M. Dunlavy, T. G. Kolda, and M. Mørup. Scalable tensor factorizations for incomplete data. Chemometrics and Intelligent Laboratory Systems, 106(1):41–56, 2011.

4. E. Acar, T. G. Kolda, and D. M. Dunlavy. All-at-once optimization for coupled matrix and tensor factorizations. In KDD Workshop on Mining and Learning with Graphs, 1105.3422, 2011.

5. E. Acar, E. E. Papalexakis, G. Gurdeniz, M. A. Rasmussen, A. J. Lawaetz, M. Nilsson, and R. Bro. Structure-revealing data fusion. BMC Bioinformatics, 15:239, 2014.

6. E. Acar, C. Schenker, Y. Levin-Schwartz, V. Calhoun, and Tulay Adali. Unraveling diagnostic biomarkers of schizophrenia through structure-revealing fusion of multi-modal neuroimaging data. Frontiers in Neuroscience, 13(416), 2019.

7. E. Acar and B. Yener. Unsupervised multiway data analysis: A literature survey. IEEE Transactions on Knowledge and Data Engineering, 21(1):6–20, 2009.

8. T. Adali, F. Kantar, M. A. B. S. Akhonda, S. Strother, V. D. Calhoun, and E. Acar. Reproducibility in matrix and tensor decompositions: Focus on model match, interpretability, and uniqueness. IEEE Signal Processing Magazine, 39(4):8–24, 2022.

9. V. Babbar, Z. Guo, and C. Rudin. What is different between these datasets? 2403.05652, 2024.

10. M. Babu and M. Snyder. Multi-omics profiling for health. Moleculer and Cellular Proteomics, 22(6), 2023.

11. B. W. Bader, T. G. Kolda, et al. Matlab tensor toolbox, version 3.1. https://www.tensortoolbox.org.

12. K. M. Bermingham, M. Mazidi, P. W. Franks, T. Maher, A. M. Valdes, I. Linenberg, J. Wolf, G. Hadjigeorgiou, T. D. Spector, C. Menni, J. M. Ordovas, S. E. Berry, and W. L. Hall. Characterisation of fasting and postprandial NMR metabolites: Insights from the ZOE PREDICT 1 study. Nutrients, 15(11):2638, 2023.

13. S. E. Berry, A. M. Valdes, D. A. Drew, F. Asnicar, M. Mazidi, J. Wolf, J. Capdevila, G. Hadjigeorgiou, R. Davies, H. Al Khatib, et al. Human postprandial responses to food and potential for precision nutrition. Nature Medicine, 26(6):964–973, 2020.

14. H. Bisgaard. The Copenhagen Prospective Study on Asthma in Childhood (COPSAC): design, rationale, and baseline data from a longitudinal birth cohort study. Annals of Allergy, Asthma & Immunology, 93(4):381–389, 2004.

15. B. J. Blaise, G. D. S. Correia, G. A. Haggart, I. Surowiec,C. Sands, M. R. Lewis, J. T. M. Pearce, J. Trygg, J. K. Nicholson, E. Holmes, and T. M. D. Ebbels. Statistical analysis in metabolic phenotyping. Nature Protocols, 16(9):4299–4326, 2021.

16. R. Bro and A. K. Smilde. Centering and scaling in component analysis. Journal of Chemometrics, 17(1):16– 33, 2003.

17. J. D. Carroll and J. J. Chang. Analysis of individual differences in multidimensional scaling via an N-way generalization of ‘Eckart-Young’ decomposition. Psychometrika, 35:283–319, 1970.

18. D. M. Dunlavy, T. G. Kolda, and E. Acar. Poblano v1.0: A Matlab toolbox for gradient-based optimization. Technical report, Sandia National Laboratories, 2010.

19. S. Fujita, Y. Karasawa, K. Hironaka, Y. Taguchi, and S. Kuroda. Features extracted using tensor decomposition reflect the biological features of the temporal patterns of human blood multimodal metabolome. PLoS ONE, 18(2):e0281594, 2023.

20. R. A. Harshman. Foundations of the PARAFAC procedure: Models and conditions for an ‘explanatory’ multi-modal factor analysis. UCLA working papers in phonetics, 16:1– 84, 1970.

21. F. L. Hitchcock. The expression of a tensor or a polyadic as a sum of products. Journal of Mathematics and Physics, 6(1):164–189, 1927.

22. B. Hunyadi, P. Dupont, W. Van Paesschen, and S. Van Huffel. Tensor decompositions and data fusion in epileptic electroencephalography and functional magnetic resonance imaging data. WIREs Data Mining and Knowledge Discovery, 7(1):e1197, 2017.

23. C. I. Kanatsoulis, X. Fu, N. D. Sidiropoulos, and W.-K. Ma. Hyperspectral super-resolution: A coupled tensor factorization approach. IEEE Transactions on Signal Processing, 66(24):6503–6517, 2018.

24. S. A. Khan, E. Leppaaho, and S. Kaski. Bayesian multi-tensor factorization. Machine Learning, 105:233–253, 2016.

25. T. G. Kolda and B. W. Bader. Tensor decompositions and applications. SIAM Review, 51(3):455–500, 2009.

26. H. Kurata. Virtual metabolic human dynamic model for pathological analysis and therapy design for diabetes. iScience, 24(2):102101, 2021.

27. G. Lépine, M. Tremblay-Franco, S. Bouder, L. Dimina, H. Fouillet, F. Mariotti, and S. Polakof. Investigating the postprandial metabolome after challenge tests to assess metabolic flexibility and dysregulations associated with cardiometabolic diseases. Nutrients, 14(3):472, 2022.

28. L. Li, H. Hoefsloot, A. A. Graaf, E. Acar, and A. K. Smilde. Exploring dynamic metabolomics data with multiway data analysis: A simulation study. BMC Bioinformatics, 23(31), 2022.

29. L. Li, S. Yan, B. M. Bakker, H. Hoefsloot, B. Chawes, D. Horner, M. A. Rasmussen, A. K. Smilde, and E. Acar. Analyzing postprandial metabolomics data using multiway models: A simulation study. BMC Bioinformatics, 25(94), 2024.

30. L. Li, S. Yan, D. Horner, M. A. Rasmussen, A. K. Smilde, and E. Acar. Revealing static and dynamic biomarkers from postprandial metabolomics data through coupled matrix and tensor factorizations. Metabolomics, 20:86, 2024.

31. C. Martino, L. Shenhav, C. Marotz, G. Armstrong, D. McDonald, Y. Vázquez-Baeza, J. T. Morton, L. Jiang, M. G. Dominguez-Bello, A. D. Swafford, E. Halperin, and R. Knight. Context-aware dimensionality reduction deconvolutes gut microbial community dynamics. Nature Biotechnology, 39:165–168, 2021.

32. E. Müllner, H. E. Röhnisch, C. Von Brömssen, and A. A. Moazzami. Metabolomics analysis reveals altered metabolites in lean compared with obese adolescents and additional metabolic shifts associated with hyperinsulinaemia and insulin resistance in obese adolescents: A cross-sectional study. Metabolomics, 17(1):1–13, 2021.

33. D. J. Panyard, B. Yu, and M. P. Snyder. The metabolomics of human aging: Advances, challenges, and opportunities. Science Advances, 8(42):eadd6155, 2022.

34. E. E. Papalexakis, C. Faloutsos, and N. D. Sidiropoulos. Tensors for data mining and data fusion: Models, applications, and scalable algorithms. ACM Transactions on Intelligent Systems and Technology, 8(2), 2016.

35. L. Pellis, M. J. van Erk, B. van Ommen, G. C. M. Bakker, H. F. J. Hendriks, N. H. P. Cnubben, R. Kleemann, E. P. van Someren, I. Bobeldijk, C. M. Rubingh, and S. Wopereis. Plasma metabolomics and proteomics profiling after a postprandial challenge reveal subtle diet effects on human metabolic status. Metabolomics, 8:347–359, 2012.

36. N. D. Price, A. T. Magis, J. C. Earls, G. Glusman, R. Levy, C. Lausted, D. T. McDonald, U. Kusebauch, C. L. Moss, Y. Zhou, S. Qin, R. L. Moritz, K. Brogaard, G. S. Omenn, J. C. Lovejoy, and L. Hood. A wellness study of 108 individuals using personal, dense, dynamic data clouds. Nature Biotechnology, 35:747–756, 2017.

37. Y. J. W. Rozendaal, Y. Wang, Y. Paalvast, L. L. Tambyrajah, Z. Li, K. Willems van Dijk, P. C. N. Rensen, J. A. Kuivenhoven, A. K. Groen, P. A. J. Hilbers, and N. A. W. van Riel. In vivo and in silico dynamics of the development of metabolic syndrome. PLOS Computational Biology, 14(6):e1006145, 2018.

38. C. Schenker, J. E. Cohen, and E. Acar. A flexible optimization framework for regularized matrix-tensor factorizations with linear couplings. IEEE Journal of Selected Topics in Signal Processing, 15(3):506–521, 2021.

39. X. Shen, R. Kellogg, D. J. Panyard, N. Bararpour, K. E. Castillo, B. Lee-McMullen, A. Delfarah, J. Ubellacker, S. Ahadi, Y. Rosenberg-Hasson, and A. Ganz. Multi-omics microsampling for the profiling of lifestyle-associated changes in health. Nature Biomedical Engineering, 2023.

40. U. Simsekli, B. Ermis, A. T. Cemgil, and E. Acar. Optimal weight learning for coupled tensor factorization with mixed divergences. In EUSIPCO’13: Proceedings of 21st European Signal Processing Conference, pages 1–5, 2013.

41. V. Skantze, M. Wallman, A. S. Sandberg, R. Landberg, M. Jirstrand, and C. Brunius. Identification of metabotypes in complex biological data using tensor decomposition. Chemometrics and Intelligent Laboratory Systems, 233:104733, 2023.

42. A. K. Smilde, P. Geladi, and R. Bro. Multi-Way Analysis with Applications in the Chemical Sciences. Wiley, 2004.

43. M. Sørensen and L. D. De Lathauwer. Coupled canonical polyadic decompositions and (coupled) decompositions in multilinear rank-(Lr,n,Lr,n,1) terms—part I: Uniqueness. SIAM Journal on Matrix Analysis and Applications, 36(2):496–522, 2015.

44. J. H. M. Stroeve, H. Van Wietmarschen, B. H. A. Kremer, B. Van Ommen, and S. Wopereis. Phenotypic flexibility as a measure of health: The optimal nutritional stress response test. Genes & Nutrition, 10(3):1–21, 2015.

45. N. Swainston, K. Smallbone, H. Hefzi, P. D. Dobson, and J. Brewer et al. Recon 2.2: from reconstruction to model of human metabolism. Metabolomics, 12(7), 2016.

46. I. Thiele, S. Sahoo, A. Heinken, J. Hertel, L. Heirendt, M. K. Aurich, and R. M. T. Fleming. Personalized whole-body models integrate metabolism, physiology, and the gut microbiome. Molecular Systems Biology, 16(5):e8982, 2020.

47. I. Thiele, N. Swainston, R. M. T. Fleming, A. Hoppe, and S. Sahoo et al. A community-driven global reconstruction of human metabolism. Nature Biotechnology, 31(5):419–425, 2013.

48. L. von Rueden, S. Mayer, K. Beckh, B. Georgiev, S. Giesselbach, R. Heese, B. Kirsch, J. Pfrommer, A. Pick, R. Ramamurthy, M. Walczak, J. Garcke, C. Bauckhage, and J. Schuecker. Informed machine learning – a taxonomy and survey of integrating prior knowledge into learning systems. IEEE Transactions on Knowledge and Data Engineering, 35(1):614–633, 2023.

49. T. F. Wilderjans, E. Ceulemans, I. Van Mechelen, and R. A. van den Berg. Simultaneous analysis of coupled data matrices subject to different amounts of noise. British Journal of Mathematical and Statistical Psychology, 64:277–290, 2011.

50. A. H. Williams, T. H. Kim, F. Wang, S. Vyas, S. I. Ryu, K. V. Shenoy, M. Schnitzer, T. G. Kolda, and S. Ganguli. Unsupervised discovery of demixed, low-dimensional neural dynamics across multiple timescales through tensor component analysis. Neuron, 98(6):1099–1115.e8, 2018.

51. M. K. Wojczynski, S. P. Glasser, A. Oberman, E. K. Kabagambe, P. N. Hopkins, M. Y. Tsai, R. J. Straka, J. M. Ordovas, and D. K. Arnett. High-fat meal effect on LDL, HDL, and VLDL particle size and number in the Genetics of Lipid-Lowering Drugs and Diet Network (GOLDN): An interventional study. Lipids in Health and Disease, 10(1):181, 2011.

52. S. Wopereis, J. H. M. Stroeve, A. Stafleu, G. C. M. Bakker, J. Burggraaf, M. J. van Erk, L. Pellis, R. Boessen, A. A. F. Kardinaal, and B. van Ommen. Multi-parameter comparison of a standardized mixed meal tolerance test in healthy and type 2 diabetic subjects: The PhenFlex challenge. Genes & Nutrition, 12(21):1–14, 2017.

53. S. Yan, L. Li, D. Horner, P. Ebrahimi, B. Chawes, L. O. Dragsted, M. A. Rasmussen, A. K. Smilde, and E. Acar. Characterizing human postprandial metabolic response using multiway data analysis. Metabolomics, 20:50, 2024.

