## Supplementary Material for "Longitudinal metabolomics data analysis informed by mechanistic models"

### Supplementary File

#### A Females: CP model demonstrates comparable performance relative to joint analysis

We analyze the real T0-corrected metabolomics measurements from females $\mathfrak{X}$ (152 females $\times$ 6 metabolites $\times$ 7 time points) using a 3-component CP model with Tikhonov regularization (with $\gamma = 0.01$), and jointly analyze real and simulated data using a 3-component ACMTF model.

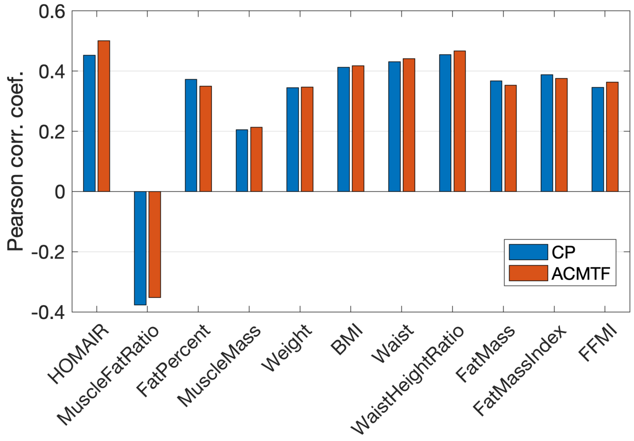


Figure S.1: Correlations between meta variables and the subject scores for the component of interest, i.e., the component that demonstrates statistically significant correlation with BMI, captured using a 3-component CP model of real data from females vs. a 3-component ACMTF model of real data from females and simulated data.

Fig S.1 shows that correlations with meta variables obtained using a CP model are comparable to the ones obtained using joint analysis. On the other hand, joint analysis of real data from males and simulated data improves the correlations as shown in Fig. 4a in the main text. To understand the different performance in males vs. females, we compare the factors extracted by the CP and ACMTF models for males and females. Fig S.2 shows that in females, the CP model extracts more similar subject components $\mathbf{a}_2$ (the pattern which shows statistically significant correlation with BMI) in CP vs. ACMTF compared to males. Taking into account of all three components, the factor match score (FMS) between CP and ACMTF factors extracted from females (FMS=0.70) is slightly larger than the FMS for males (FMS=0.63).

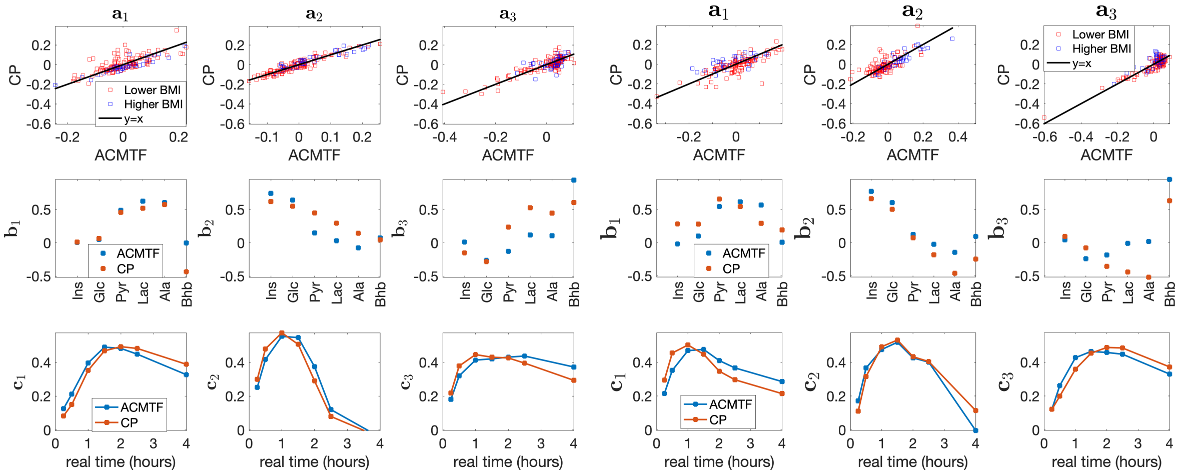


(a) Female (b) Male

Figure S.2: Comparisons of the factors extracted by 3-component CP and ACMTF models for (a) females, and (b) males.

### B Males: Selection of the number of components and regularization parameter

#### B.1 CP model of real metabolomics data

In our analysis, we observe that CP models using 2, 5, and 6 components without any regularization are degenerate [2]; therefore, we consider 3- and 4-component CP models. Fig S.3a shows the replicability of the extracted factors indicating that 95% of the FMS values (see the definition of FMS in subsection *Model Selection* in the main manuscript) using the 3-component and 4-component models are greater than 0.87 and 0.58, respectively. Therefore, we select a 3-component CP model.

In order to determine the regularization parameter, we have assessed replicability of the 3-component CP model using different regularization parameter values ($\gamma$). Fig S.3b shows that $\gamma = 0.01$ is a good choice in terms of both replicability and the model fit.

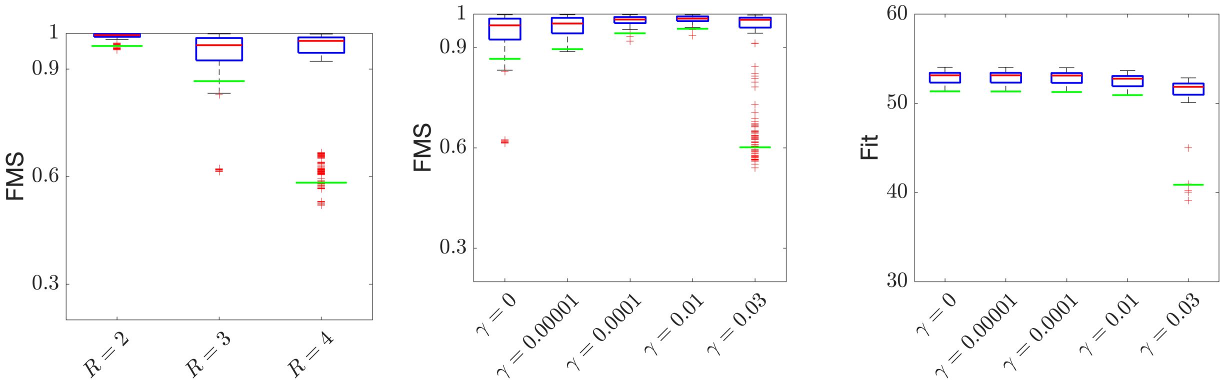


(a) Replicability–no regularization (b) Replicability and model fit using different regularization parameters

Figure S.3: Model selection for the CP model of T0-corrected data from males. (a) Replicability of the CP model without any regularization using different number of components, (b) Replicability and model fit (%) of the 3-component CP model using different regularization parameters. Green lines show that 95% of the FMS and model fit values are above that line.

#### B.2 ACMTF model of joint analysis of real and simulated metabolomics data

In our investigation, the 2-component, 3-component, and 4-component ACMTF models are replicable as shown in Fig S.4a. When we use a 4-component model, the first component in the 3-component model (i.e., $\langle \mathbf{a}_1, \mathbf{b}_1, \mathbf{c}_1, \mathbf{d}_1, \mathbf{e}_1 \rangle$ in Fig S.4b) which mainly models Pyr, Lac and Ala splits into two components in the

4-component model (i.e., $\langle \mathbf{a}_1, \mathbf{b}_1, \mathbf{c}_1, \mathbf{d}_1, \mathbf{e}_1 \rangle$ and $\langle \mathbf{a}_4, \mathbf{b}_4, \mathbf{c}_4, \mathbf{d}_4, \mathbf{e}_4 \rangle$ in Fig S.4c). Given that Pyr, Lac, and Ala exhibit similar dynamic patterns and are close in the metabolic pathway [1], it is rational to maintain their grouping within a single component. Therefore, we select the 3-component ACMTF model.

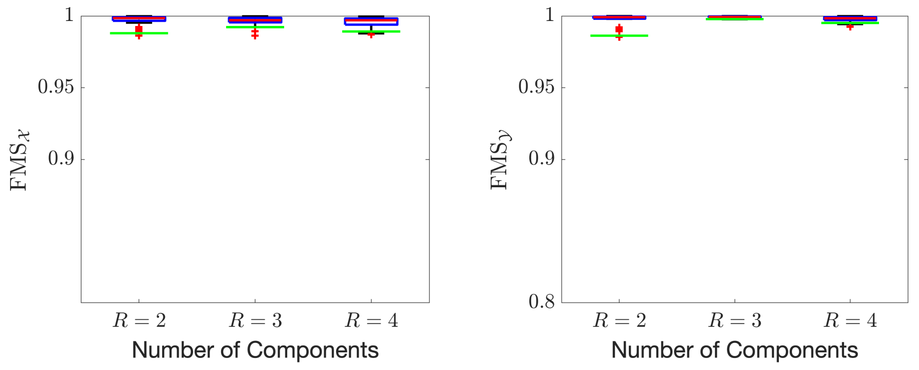


(a) Replicability

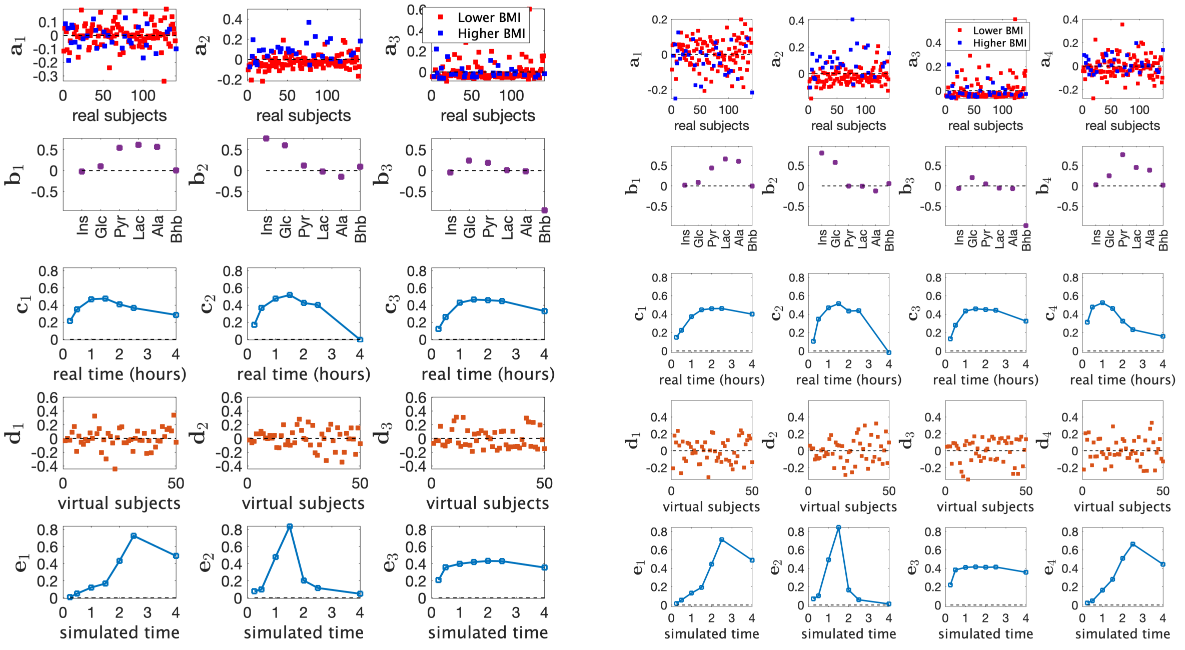


(b) (c)

Figure S.4: Model selection for the ACMTF model of T0-corrected real data (from males) and simulated data. $\langle \mathbf{a}_r, \mathbf{b}_r, \mathbf{c}_r, \mathbf{d}_r, \mathbf{e}_r \rangle$, $r = 1, 2, 3, 4$, are the components in the *subjects* (real), *metabolites* (coupled mode), *time* (real), *subjects* (virtual) and *time* (virtual) modes.

### C Time profiles of raw data

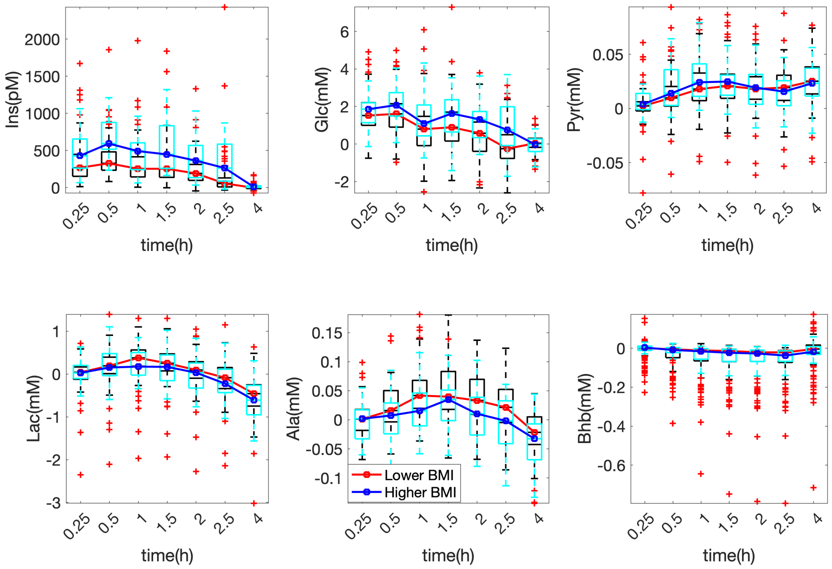


Figure S.5: Median time profiles of 111 male subjects categorized within the *Lower BMI* group vs. 30 male subjects from the *Higher BMI* group, based on T0-corrected data.

### D Conflicting prior information

#### D.1 CP model of simulated metabolomics data and construction of simulated data with conflicting patterns

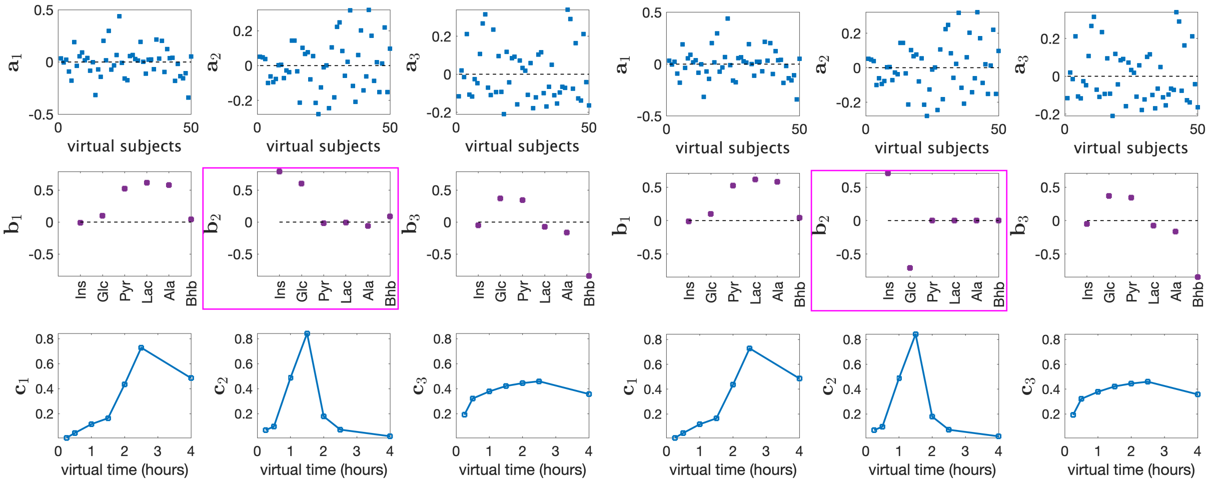


(a) Default true factors (b) Conflicting factors

Figure S.6: **(a)** True factors extracted by a 3-component CP model from the default simulated T0-corrected metabolomics data. **(b)** Modified patterns used to generate *simulated data with conflicting information*. In (a) vs. (b), only $\mathbf{b}_2$ is different. $\mathbf{b}_2$ in Fig S.6b is obtained by setting the values of Ins, Glc, Pyr, Lac, Ala, Bhb to $1, -1, 0, 0, 0, 0$ and then dividing this vector by its 2-norm.

#### D.2 Joint analysis of real data and simulated data with conflicting information

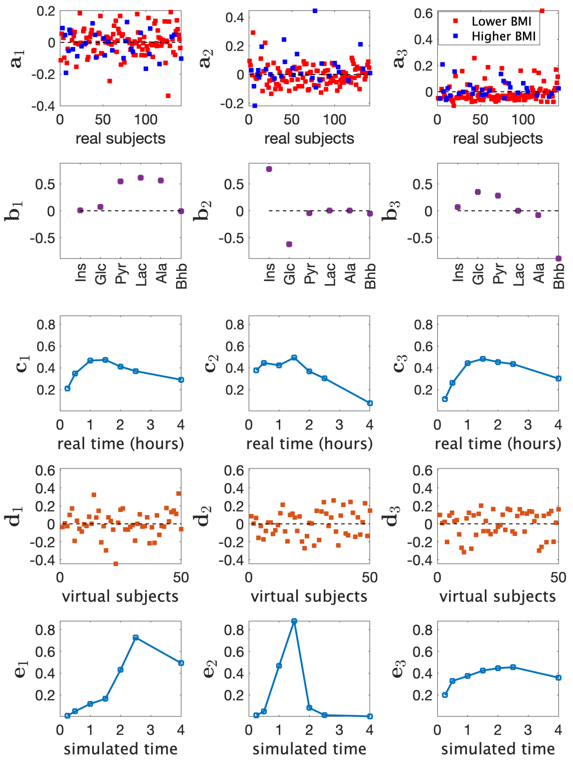


Figure S.7: Factors of the 3-component ACMTF model of T0-corrected real data from males and simulated data with conflicting patterns. $\langle \mathbf{a}_r, \mathbf{b}_r, \mathbf{c}_r, \mathbf{d}_r, \mathbf{e}_r \rangle$, $r = 1, 2, 3$, are the components in the *subjects* (real), *metabolites* (coupled), *time* (real), *subjects* (virtual) and *time* (virtual) modes.

### E Sensitivity of joint analysis to different weighting schemes

When jointly analyzing real and simulated data using an ACMTF model, we give equal importance to each data set by solving the optimization problem (1) with $\alpha = 0.5$. Here, we assess the sensitivity of the performance in terms of the correlations with the meta variables to different weighting schemes, i.e., different $\alpha$. Figure S.8a shows that correlations increase as we incorporate the simulated data in the analysis, i.e., with $\alpha > 0$ (only the CP model of simulated data) and $\alpha < 1$ (only the CP model of real data). From $\alpha = 0.8$ to $\alpha = 0.1$, the correlations show minor increase. Figure S.8b shows the factor match scores between factors of models with different weights and the factors obtained using equal weights (i.e., $\alpha = 0.5$). We observe that models with $\alpha$ varying from 0.6 to 0.1 extract similar patterns, with FMS values over 0.95. Overall, we observe that unless we are close to the extremes, the model is not very sensitive to the weight selection (based on the weights considered here). It is possible to achieve higher performance but that requires a rigorous validation process in order not to overfit the data - which is a topic of future research.

$$\begin{aligned} \min_{\boldsymbol{\lambda},\boldsymbol{\sigma},\mathbf{A},\mathbf{B},\mathbf{C},\mathbf{D},\mathbf{E}} \quad & \alpha \, \| \, \mathcal{X} - [\![ \boldsymbol{\lambda}; \mathbf{A}, \mathbf{B}, \mathbf{C} ]\!] \, \|^2 + (1-\alpha) \, \| \, \mathcal{Y} - [\![ \boldsymbol{\sigma}; \mathbf{D}, \mathbf{B}, \mathbf{E} ]\!] \, \|^2 \\ & + \beta \, \| \, \boldsymbol{\lambda} \, \|_1 + \beta \, \| \, \boldsymbol{\sigma} \, \|_1 \,, \qquad (1) \\ \text{s.t.} \; \| \, \mathbf{a}_r \, \| = \| \, \mathbf{b}_r \, \| = \| \, \mathbf{c}_r \, \| = \| \, \mathbf{d}_r \, \| = \| \, \mathbf{e}_r \, \| = 1, r = 1, \cdots, R. \end{aligned}$$

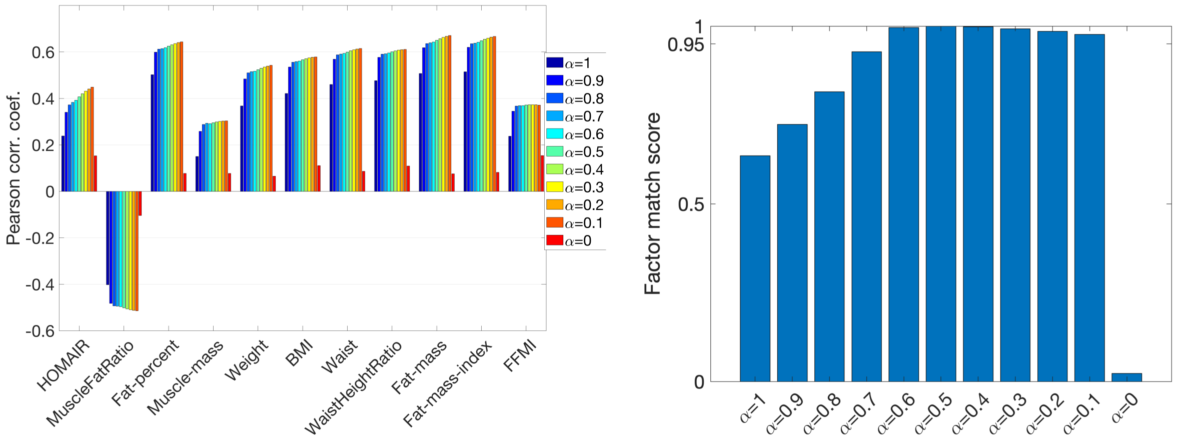


(a)

(b)

Figure S.8: Sensitivity of ACMTF models to different weighting schemes in (1). **(a)** Correlations between the subject scores and meta variables for the factor that gives the strongest correlations, **(b)** FMS between factors extracted by an ACMTF model using different weights and those obtained with equal weights (i.e., $\alpha = 0.5$).

### References

[1] L. Li, S. Yan, B. M. Bakker, H. Hoefsloot, B. Chawes, D. Horner, M. A. Rasmussen, A. K. Smilde, and E. Acar. Analyzing postprandial metabolomics data using multiway models: A simulation study. *BMC Bioinformatics*, 25(94), 2024.

[2] A. Stegeman. Degeneracy in Candecomp/Parafac and Indscal explained for several three-sliced arrays with a two-valued typical rank. *Psychometrika*, 72(4):601–619, 2007.
